# Collaborative network analysis for the interpretation of transcriptomics data in rare diseases, an application to Huntington’s disease

**DOI:** 10.1101/2023.07.22.550153

**Authors:** Ozan Ozisik, Nazli Sila Kara, Tooba Abbassi-Daloii, Morgane Térézol, Núria Queralt-Rosinach, Annika Jacobsen, Osman Ugur Sezerman, Marco Roos, Chris T. Evelo, Anaïs Baudot, Friederike Ehrhart, Eleni Mina

## Abstract

**Background:** Rare diseases may affect the quality of life of patients and in some cases be life-threatening. Therapeutic opportunities are often limited, in part because of the lack of understanding of the molecular mechanisms that can cause disease. This can be ascribed to the low prevalence of rare diseases and therefore the lower sample sizes available for research. A way to overcome this is to integrate experimental rare disease data with prior knowledge using network-based methods. Taking this one step further, we hypothesized that combining and analyzing the results from multiple network-based methods could provide data-driven hypotheses of pathogenicity mechanisms from multiple perspectives.

**Results:** We analyzed a Huntington’s disease (HD) transcriptomics dataset using six network-based methods in a collaborative way. These methods either inherently reported enriched annotation terms or their results were fed into enrichment analyses. The resulting significantly enriched Reactome pathways were then summarized using the ontological hierarchy which allowed the integration and interpretation of outputs from multiple methods. Among the resulting enriched pathways, there are pathways that have been shown previously to be involved in HD and pathways whose direct contribution to disease pathogenesis remains unclear and requires further investigation.

**Conclusions:** In summary, our study shows that collaborative network analysis approaches are well-suited to study rare diseases, as they provide hypotheses for pathogenic mechanisms from multiple perspectives. Applying different methods to the same case study can uncover different disease mechanisms that would not be apparent with the application of a single method.

## Introduction

Rare diseases are defined as diseases with low prevalence. The European Commission, for instance, considers a disease to be rare when it affects fewer than 1 person in 2,000 [1]. A recent study by Orphanet identified 6,172 unique rare diseases [2] that can be life-threatening or severely affect the quality of life of patients. Therapies are only available for a very small number of these diseases [3, 4]. Therefore, there is an immediate need to accelerate the study of these diseases, and thereby assist in the development of new therapies. However, investigating rare diseases is challenged by sample scarcity, clinical and genetic variability, and data unavailability or deficiency. Systems biology is a strategy that is used to overcome these challenges by providing systems-level insights for rare disease research. In this direction, network-based computational methods combine experimental data (e.g., omics data) with prior knowledge available in databases and ontologies. These methods can translate the sparse, rare disease-specific information into a systems-based understanding of disease pathology. This can accelerate research in rare disease and identify, for instance, deregulated pathways, biological processes and drug targets.

A first set of network methods aims to aid the interpretation of large-scale molecular data (typically transcriptomics) by adding interaction information. GeneMania [5] and STRING [6] are two tools that fetch and visualize the physical and functional interactions between genes or proteins of interest; they also allow including additional interacting proteins to the networks and performing enrichment analysis. These tools are available as both web applications and Cytoscape [7] apps. CyTargetLinker is another Cytoscape app that can extend a given list of genes or proteins using a linkset that contains interaction partners information such as gene-pathway relations or miRNA target relations. PathVisio [8] is a standalone tool that allows visualizing and analyzing genes of interest with the pathway information from WikiPathways [9]. EnrichNet [10] is a tool that uses both large-scale interaction network data and annotation data to provide enrichment for gene sets of interest.

Another set of network methods has the objective to integrate experimental data with biological networks in order to identify subnetworks of interest, also known as active modules [11]. Exploring all potential subnetworks is a complex task from a computational point of view, and methods based on different algorithms have been developed to overcome this challenge [11]. For instance, jActiveModules [12] is a method based on simulated annealing that has been widely used. More recent solutions include MOGAMUN [13] that is based on a multi-objective genetic algorithm, pathfindR [14], which by default uses greedy search, and DOMINO [15], which applies multiple steps of network partitioning.

Finally, another set of network methods uses statistical methods to infer interactions between molecules from the expression data, creating for instance co-expression networks. Co-expression networks are constructed by computing the pairwise correlation between expression profiles of molecules in different conditions. A popular method in this field is Weighted Gene Co-expression Network Analysis (WGCNA) [16], which targets a scale-free topology in the resulting network. Weighted topological overlap (wTO) [17] is another method to build co-expression networks which considers both positive and negative correlations. The inferred networks are then piped with subsequent network analysis tools, such as module detection or co-expression differential network analysis [18, 19, 20]. A recently developed tool, MODifier, combines multiple methods for network inference and module detection [21] to generate more robust modules.

Different network-based methods are expected to extract different information from the data. We hypothesize that the combination of multiple network-based methods can strengthen and enhance our understanding of the mechanisms involved in rare disease pathogenesis. Indeed, methods with different viewpoints, assumptions and prior knowledge can provide both supportive and complementary findings that lead to more robust and enhanced results when aggregated.

In accordance with this hypothesis, we followed a collaborative approach in which we selected a diverse set of methods representative of the different network-based methodological categories introduced above. These methods are WGCNA, wTO-CoDiNA, PathVisio/Cytoscape/CyTargetLinker, EnrichNet, pathfindR and MOGAMUN. We analyzed the Huntington’s Disease (HD) gene expression dataset described in the study by Labadorf et al. [22], as a case study of a rare disease. HD is a dominantly inherited neurodegenerative disorder with a prevalence of 10.6-13.7 individuals per 100,000 in Western populations [23]. The cause of the disease, the mutated huntingtin protein, was discovered in 1993 [24]. However, symptomatic management is the current way of treatment and we still lack disease-modifying treatments [25]. The huntingtin gene (*HTT*) mutation corresponds to a repeat expansion of the CAG codon, which translates to a polyglutamine expansion (polyQ) in the encoded huntingtin protein [24]. Clinically, an extensive brain degeneration is primarily responsible for the most common symptoms that are evident from the early disease stages [25]. The symptoms gradually worsen over time, with death occurring approximately 15–20 years after disease onset [26].

In this project, we demonstrate that employing various network analysis methods increases the reliability of common results unveiled by different methods and provides new valuable insights that contribute to novel perspectives and a deeper understanding of diseases. Collaborative network analysis is a strategy that can be beneficial for rare disease research where the lack of data and knowledge is an important barrier.

## Results

We applied six network-based methods that represent the three method categories described in the introduction: interaction mapping, active module identification and network inference.

PathVisio/Cytoscape/CyTargetLinker and EnrichNet which use interaction information for biological interpretation; pathfindR and MOGAMUN which identify subnetworks of interest by integrating experimental data with biological networks; and finally WGCNA and wTO-CoDiNA which infer gene interactions with statistical inference. These methods are applied to the Huntington’s disease (HD) gene expression dataset provided by Labadorf et al. [22] (Figure 1, see Methods section for details on the data and the methods). Depending on the input requirements of each method, we used either the normalized count data or the differential expression analysis data (both are retrieved from the original study and available through NCBI Gene Expression Omnibus, accession number GSE64810). We collected and assessed the outputs of all the methods at the level of enriched pathways, using the Reactome database [27]. Two of the methods, i.e., EnrichNet and pathfindR, have incorporated functions for enrichment analysis. The other four methods do not have intrinsic enrichment analysis function. In these cases, we used g:Profiler [28]. Since the Reactome database is hierarchically organized, the enrichment results included both general terms and more specific subterms that are descendents of the general terms. As a final step, we used an enrichment analysis filtering tool, orsum [29], to summarize and integrate the results obtained from different methods. orsum selects more significant (higher ranked) general terms as representatives for their less significant (lower ranked) subterms.

**Figure 1.**
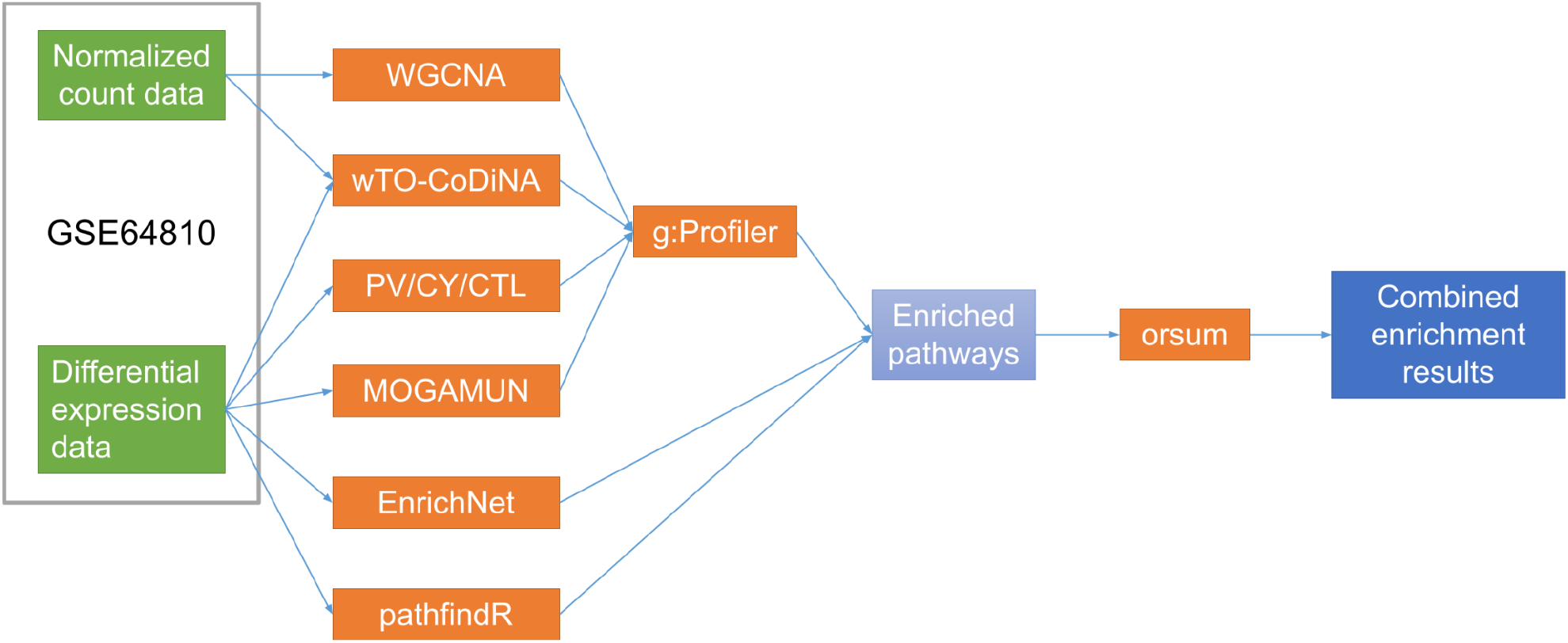
Collaborative network analysis workflow.

Our collaborative approach resulted in a total of 649 enriched Reactome terms from the six network-based methods. The application of orsum filtering led to 109 representative Reactome terms (Figure 2). Please note that, while orsum integrated the results from different methods and provided a summarized view, it is still important to examine the filtered out subterms, as they differ between methods and can reflect more detailed biological mechanisms. Therefore, we discuss both the representative Reactome terms and their associated subterms that emerged as significant in our analysis.

**Figure 2.**
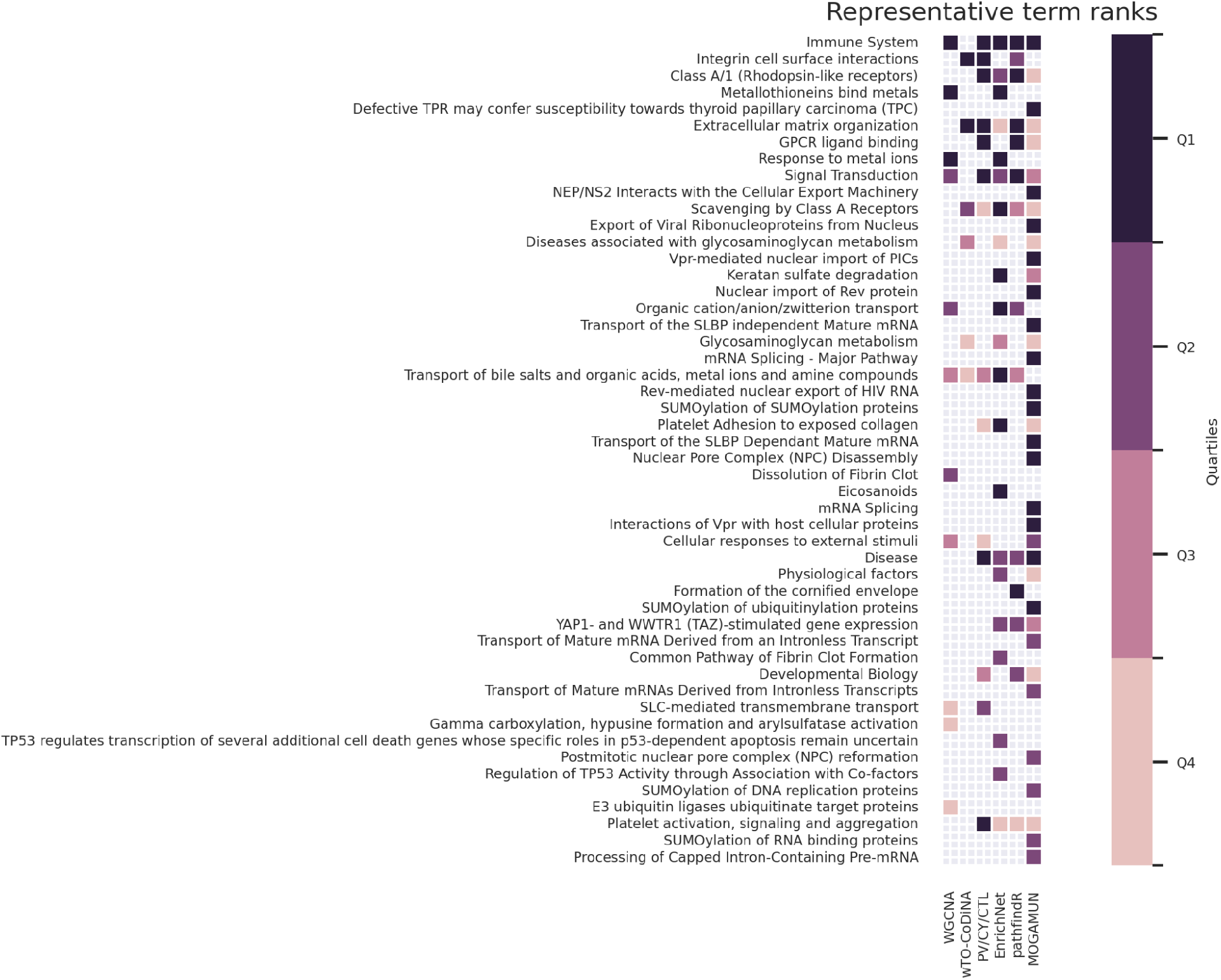
Result of Reactome pathway enrichments from the six network-based methods, summarized by orsum. The top 50 terms are presented.

“Immune System” and its subterms are ranked at the top of the results. “Innate Immune System”, “Signaling by Interleukins”, “Cytokine Signaling in Immune system”, “Neutrophil degranulation” are some of the top ranking terms found by most of the network-based methods. Immune system activation is well known in HD and both innate and adaptive immune systems are thought to be playing a role in HD pathology [30, 31]. In our study, all methods except wTO-CoDiNA have identified enriched Reactome terms related to the immune system.

“Signal Transduction” and/or its subterms are significantly enriched according to all the methods, except wTO-CoDiNA. “Signal Transduction” is a generic term annotating more than 2,500 protein-coding genes and many different subterms were summarized by it. “Signaling by TGFB family members” and “Signaling by Receptor Tyrosine Kinases” are among the significant subterms. Alterations in numerous signal transduction pathways are well known in HD, for example elevation of TGF-beta signaling component [32], disruption of Signaling by EGFR [33] and alterations of MAPK and ERK signaling pathways [34].

“Extracellular matrix organization” and/or its subterms are significantly enriched according to all the methods, except WGCNA. Extracellular matrix (ECM) proteins are altered during regular aging and nowadays it is clear that they are also altered during disease. In particular, changes in the expression of ECM proteins have been reported for three neurodegenerative diseases including HD [35]. While a clear direct link between HD and ECM changes has not been reported in humans, a recent study in HD mouse model reported that microglia elimination leads to prevention of extracellular matrix changes [36].

“Transport of bile salts and organic acids, metal ions and amine compounds”, a process mediated by solute carriers (SLCs), was identified by all methods, except MOGAMUN. SLCs play an important role in various neurodegenerative diseases [37] and they were found to be dysregulated in multiple HD studies [38, 39, 40]. Hence, SLCs have been suggested as important therapeutic targets [41]; a well known example is tetrabenazine, an inhibitor of SLC18A2 that is used to treat chorea in HD [42].

“Platelet activation, signaling and aggregation” was also identified by all the methods except WGCNA and wTO-CoDiNA. Subterms of this pathway such as “Platelet degranulation”, “Response to elevated platelet cytosolic Ca2+” and “GP1b-IX-V activation signaling” are among the enriched terms. Platelet activation, signaling and aggregation are the three major processes for the establishment of hemostasis [43]. Platelet abnormalities have been detected in HD, which in turn promote blood-brain barrier permeability [44].

“Developmental Biology” was identified by PV/CY/CTL, pathfindR and MOGAMUN. This coincides with one of the key findings of Labadorf et al. [22] where developmental genes were identified as highly overrepresented. The significant subterms of this general term include the terms associated with neuronal system development (identified by PV/CY/CTL and pathfindR) and other developmental biology pathways such as “Signaling by NODAL”, “Transcriptional regulation of pluripotent stem cells” and “Myogenesis” (MOGAMUN). HD, by its nature as a neurodegenerative disorder, has been linked with neuronal system development [26].

“Metallothioneins bind metals” was ranked very high by two of our methods, EnrichNet and WGCNA. Metallothioneins was one of the most enriched terms in the original publication by Labadorf et al. [22]. In another study, it was also shown that metallothioneins can protect against polyQ toxicity in two cellular HD model systems and therefore were proposed as a candidate therapeutic target for HD [45].

“Defective TPR may confer susceptibility towards thyroid papillary carcinoma (TPC)”, “Nuclear Pore Complex (NPC) Disassembly”, “Postmitotic nuclear pore complex (NPC) reformation” are a few examples of the pathways that ranked very high and were identified by a single method, MOGAMUN. There is increasing evidence that mutant huntingtin disrupts the nucleocytoplasmic transport and the NPC function. New studies indeed suggest that defects in the NPC might be an important pathogenic mechanism and could possibly become a new therapeutic avenue for HD [46, 47, 48, 49].

## Discussion

The increasing demands and complexity of biomedical research requires collaborative interdisciplinary research. We hypothesized that, in the investigation of rare disease mechanisms, the combination of multiple network-based methods can lead to more robust and enhanced results when aggregated. Different network-based methods provide different viewpoints as they are implemented under different assumptions and they use different sources of prior knowledge.

In this study, we applied six network-based methods, namely WGCNA, wTO-CoDiNA, PathVisio/Cytoscape/CyTargetLinker, EnrichNet, MOGAMUN and pathfindR, to the same use case and dataset. Our use case is a rare neurodegenerative disorder, Huntington’s disease (HD), and the dataset is provided by Labadorf et al. [22]. In the case of HD, the scarcity of disease tissue and the limited number of patients hinders the production of relevant datasets. Thus, collaborative network analysis that builds on the synergy between human expertise and application of different analytical methods becomes essential for facilitating knowledge discovery in HD, as well as in other rare diseases.

Overall, we observe that there is an overlap between the results of all the methods, as they were able to detect the main disease signal, adding strong evidence on what was also reported by Labadorf et al. [22]. Increased inflammation and altered developmental processes were key findings of the original publication and were also reported in a high ranking order by our methods. In addition, signal transduction has largely been implicated in HD and was identified by the majority of our methods. Platelet activation, signaling and aggregation and related functions were also identified by the majority of our methods, although the direct clinical contribution to HD still remains elusive. This result points out that further research may reveal pathogenic insights on the impact of mutant huntingtin on regular platelet function. There are also pathways relevant for HD which were identified by only one or two methods, e.g., “Nuclear pore complex” and “Metallothioneins bind metals” related pathways [46, 45]. Metallothioneins were one of the most enriched clusters in the original publication by Labadorf et al. [22], and interestingly were only picked by WGCNA and EnrichNet. Alterations of metal homeostasis have been consistently reported in HD. Elevated metals in post-mortem HD but also in disease models altogether, point out that alterations in metal biology is having an impact in HD pathology [50, 51, 52]. Nuclear pore complex (NPC) has been implicated in several neurodegenerative diseases [53] by mutations in certain NPC proteins that cause different neurodegenerative phenotypes. Similarly, in HD aberrant NPC function, especially regulation of nucleocytoplasmic transport is linked to the pathogenesis of HD [46]. Although both these processes have not been extensively described in the context of HD, further study might decipher their exact role in HD pathogenesis and provide new targets for therapeutic interventions.

Biological network analysis may partially depend on the applied methods and the prior knowledge on molecular mechanisms and pathways, used to support the analysis. Each algorithm has the potential to capture different aspects of biology as they are developed under different assumptions. We hypothesize that this is the reason why some processes were detected by some algorithms but not by others. For example, MOGAMUN integrates several layers of information (PPI, pathways, molecular complexes, co-expression) with the gene expression data, which led to the identification of multiple pathways that were not identified by the other algorithms. wTO-CoDiNA focuses on the differences between case and control networks and takes into account the gene pairs coexpressed and connected only in the case but lacking this co-expression relation in the control. Nevertheless, linking the discovered unique processes to the specifics of each algorithm is beyond the scope of this study.

One limitation of our collaborative approach is that we did not consider an extensive list of network analysis methods nor perform a benchmark for selecting the methods. However, we considered the heterogeneity of the network methods when including them in this study. We indeed selected network methods that were representative of each broad category discussed earlier in the introduction.

To the best of our knowledge, this is the first study applying different network-based analysis methods in a collaborative way for a rare disease. The results show that our approach can identify disease-related biological pathways that would be missed if a single network-based method was used. Different methods can provide complementary insights at once and also contribute to an increased validity of the results that were discovered in common. Importantly, it opens new avenues for HD research to explore novel hypotheses.

## Conclusion

In this study, we followed a collaborative approach in which we applied six network-based methods to Huntington’s disease transcriptomics data. The goal was to discover disease related mechanisms and investigate the benefits of using multiple methods in terms of complementariness and validation.

Collaborative network analysis has mainly two advantages over the solo analyses: First, the use of multiple methods proposes different perspectives on the same data due to the various specifics of each algorithm. This helps in the discovery of hidden disease mechanisms for rare disorders, which is an advantage for the field, due to the limitations of rare disease studies. Second, detection of the same or similar pathways and biological processes by multiple methods increases the validity of the results and creates stronger candidate mechanisms in explaining rare diseases. Collaborative network analysis introduces a broader perspective of the disorder under scope, and we recommend its use especially in the case of rare disease research, where data and knowledge are scarce.

## Methods

### Data source

The RNAseq transcriptomics dataset used as input for all the network methods applied in this study was generated by Labadorf et al. [22] and includes 69 samples: 20 samples from Huntington’s disease patients and 49 from healthy controls. The samples were collected from human post-mortem brain tissue and subjected to RNA sequencing with the Illumina HiSeq 2000 platform. The normalized count data and the DESeq2 differential expression analysis [54] data are available in GEO under accession number GSE64810. WGCNA used the normalized count data, wTO-CoDiNA used both the normalized count data and the differential expression analysis data, and all the remaining methods used the differential expression analysis data.

In the enrichment analysis step, we used the Reactome pathway annotations [27]. For the network methods that do not have incorporated functions for enrichment analysis (i.e., MOGAMUN, WGCNA, wTO-CoDiNA and PathVisio/Cytoscape/CyTargetLinker), we used g:Profiler version e101_eg48_p14_60e968a [28] with the default parameters (multiple testing correction method is “g:SCS” and the user threshold for enrichment is 0.05). For the methods with incorporated enrichment analysis functions (i.e., EnrichNet and pathfindR), we used the same Reactome pathway dataset as the one used by g:Profiler (Reactome, annotations: BioMart, classes: 2020-10-12).

### Network-based analysis methods

We selected six network-based analysis methods that represent the different categories described in the Introduction: WGCNA and wTO-CoDiNA, which infer gene interactions with statistical inference; PathVisio/Cytoscape/CyTargetLinker and EnrichNet, which use interaction information for biological interpretation; and finally pathfindR and MOGAMUN, which identify subnetworks of interest by integrating experimental data such as transcriptomics data with biological networks. The methods are detailed below and the code used for each method is listed in the Availability of data and materials section.

### WGCNA

Weighted gene co-expression network analysis (WGCNA) [16, 55] is an algorithm that constructs a gene network and groups genes with similar expression profiles in the same subnetwork (module). The identification of gene-gene interactions and modules is based on pairwise correlations between those genes. In addition, for each module, an eigenvector is computed and correlated to the disease status to identify modules associated with the disease.

As input to the WGCNA R package (v. 1.71), we used the HD normalized count data. In order to calibrate the parameters of the WGCNA network, we used the approach presented by Abbassi-Daloii et. al (2020) [56]. Briefly, this approach uses prior knowledge of gene interactions from a pathway database to select the most optimal set of WGCNA parameters. Using this approach, we tested various combinations of WGCNA settings for power, minClusterSize, deepSplit and CutHeight by a full parameter sweep. To assess these different settings, we used the knowledge network obtained from the Reactome database using the g:Profiler R package version 0.2.0. Based on this assessment, we assigned all possible pairs of genes to 4 different groups: (1) in the same WGCNA module and Reactome pathway, (2) in the same WGCNA module but not in the same Reactome pathway, (3) not in the same WGCNA module but in the same Reactome pathway, and (4) neither in the same WGCNA module nor in the same Reactome pathway. We calculated the enrichment factor ((No. pairs in group 1 * No. pairs in group 4) / (No. pairs in group 2 * No. pairs in group 3)) and selected the optimal set of parameters with the highest enrichment factor to construct the weighted gene co-expression network. The selected parameters for WGCNA were power: 14, MinModuleSize: 15, deepSplit: 4, Cut Height: 0.1. We used g:Profiler to annotate these modules with Reactome.

### wTO-CoDiNA

wTO-CoDiNA is a method that builds co-expression networks by using weighted topological overlap and gene expression data and compares the obtained networks to detect their similarities and differences. This method uses two networks (case and control), compares these networks and shows which coexpression patterns are common for both networks, or which connections are unique to only one network. Through this comparison, the method illustrates the change of co-expression patterns between gene pairs and enables the biological interpretation of the differences between case and control groups.

In this method, we used normalized count data and the differential expression analysis data, both extracted from GEO under accession number GSE64810. The differential expression data were filtered to keep the genes with adjusted p-value ≤ 0.05 and |log2FC| > 1. After this filtering, we retrieved the gene list to be used in order to create count matrices from normalized count data. Two count matrices were created for case and control groups. The expression values of differentially expressed genes were used to construct case and control networks. Two co-expression networks were built via the count matrices, one for cases and one for controls. The R package wTO (version: 1.6.3) [17] that uses weighted topological overlap measure was used in the R environment. Non-correlated gene pairs (wTO.wTO_abs = 0.00) and correlations with adjusted p-value > 0.05 were filtered out to obtain final case and control networks. Network comparison was completed with Co-expression Differential Network Analysis (CoDiNA), by applying the R package CoDiNA (version: 1.1.2) [20]. The output of CoDiNA shows the common (alpha), differentiating (beta) and unique (gamma) connections between genes among case and control networks. In order to focus on the pathways and biological processes that may be responsible for the disease, we focused on the gene pairs connected only in the case group but not in the control group. These gene pairs were linked by “gamma links”, which show a disease-specific connection of a gene pair. In order to select the well-defined gamma links and to eliminate weak links, only the gamma links with Score_ratio ≥ 1 and Score_center > mean(Score_center) were kept. Gene pairs connected by the remaining gamma links were submitted to g:Profiler for enrichment analysis.

### PathVisio/Cytoscape/CyTargetLinker

PathVisio/Cytoscape/CyTargetLinker is an approach that represents the integrated usage of three tools. PathVisio [8] is a tool to visualize pathways and analyze genes of interest. Cytoscape [7] is a software platform for visualizing networks and integrating them with attribute data. Finally, CyTargetLinker [57] is a Cytoscape app that can extend the given list of genes or proteins using a linkset that contains interaction partners information such as gene-pathway relations or miRNA target relations. We used PathVisio version 3.3.0 and pathway overrepresentation analysis was conducted using the WikiPathways human pathway collection (version 10.10.2020) [9] and the built-in analysis function of PathVisio. As input, we used the differential expression analysis data, respectively, the genes with a p-value ≤ 0.05 and |log2FC| > 1.

From the original 28,000 gene identifiers in the transcriptomics dataset, 10,196 were recognized by WikiPathways, indicating the annotation of the gene in at least one of the (human) pathways. Of these, 668 genes met the criterion for being differentially expressed. Pathways were considered as overrepresented if the z-score was >1.96 and there were more than 3 differentially expressed genes in the pathway found. This led to 61 overrepresented pathways. These pathways were imported to Cytoscape (version 3.8.2) using WikiPathways app (version 3.3.7) for Cytoscape and merged. The differential expression analysis data was imported to Cytoscape as a node table. This initial network was extended using CyTargetLinker (version 4.0.0) app for Cytoscape and the WikiPathways linkset created from WikiPathways release 2021-01-10 (linksets “Homo sapiens (hsa) - curated collection” and “Homo sapiens (hsa) - Reactome collection”). The linksets contain gene-pathway association information derived from WikiPathways and Reactome. A standard network analysis as an in-built function in Cytoscape was used to determine node degrees. The ten nodes with the highest degrees were selected for enrichment analysis using g:Profiler.

### EnrichNet

EnrichNet [10] is an enrichment tool that aims to improve enrichment analysis by utilizing network topology information. EnrichNet takes a list of genes of interest as input and then identifies significantly associated processes/pathways by computing an Xd-score assessing the significance of network distances between the nodes corresponding to the genes of interest and the nodes annotated for reference pathways/processes. Any network can be used as a backbone, and network distances are measured by random walk with restart.

As input genes of interest, we used the list of differentially expressed genes with adjusted p-value ≤ 0.05 and |log2FC| > 1. We ran the EnrichNet method locally in order to use the same Reactome data with the other methods. We used BioGRID [58] protein interaction network as the backbone. We selected the top 50 Reactome pathways according to their Xd-scores for further interpretation.

### pathfindR

pathfindR [14] is an R package for active module identification and enrichment analysis. In pathfindR, active modules are extracted using one of the following three heuristic methods: greedy search, genetic algorithm or simulated annealing. Following this, the nodes in each active module are used in the enrichment analysis.

In this study, we used pathfindR v1.6.1. As input, we used the list of differentially expressed genes with adjusted p-value ≤ 0.05 and |log2FC| > 1. We ran the default greedy search approach with a maximum depth of 2 (i.e., the distance of any node in the module to the node that the search started from is at most 2). We used the BioGRID protein interaction network [58]. For the enrichment analysis, we set the gene sets argument to the specific Reactome data described in the Data source section.

### MOGAMUN

MOGAMUN [13] is a multi-objective genetic algorithm that allows the identification of active modules from the integration of transcriptomics data with a multiplex biological network (i.e., a network composed of different layers of physical or functional interactions).

As input for MOGAMUN, we used the adjusted p-values of all the genes from the differential expression analysis data and a multiplex network composed of four layers of interactions. The first three layers are obtained from [59]: A protein-protein interaction (PPI) layer corresponding to the fusion of three datasets: APID (apid.dep.usal.es, [60, 61]) (Level 2, human only), Hi-Union and Lit-BM [62]; a pathway layer extracted from NDEx [63] which is built in [64] and corresponds to the human Reactome data; A layer of molecular complexes constructed from the fusion of Hu.map [65] and CORUM [66], using OmniPathR [67]. The fourth layer contains edges corresponding to correlations of expression, where Spearman correlations were calculated from RNA-seq data of 32 tissues and 45 cell lines (extracted from proteinatlas.com), and absolute correlations of at least 0.7 were selected to build the co-expression network.

MOGAMUN was run 30 times with the default parameters. We identified 22 active modules. We used g:Profiler R package (gprofiler2, version 0.2.0) for the enrichment analysis of each module and then we merged these enrichment results by selecting each unique enriched pathway with the best p-value it obtained from any module.

### Enrichment results summarization using *orsum*

orsum [29] is a tool dedicated to filtering/summarizing enrichment analysis results. It targets the lengthy results arising from the redundancy in hierarchically organized annotation databases (e.g., GO and Reactome). It enables integration and joint filtering of the enrichment results obtained by different methods.

The enrichment analysis results obtained from the six different network methods were used as input for orsum (v1.6). For the annotation data, we used the Reactome data described in the Data source section. We ran orsum with default parameters.

## Declarations

### Availability of data and materials

All the codes, data and results are available in the Zenodo repository (https://doi.org/10.5281/zenodo.8174258). Specific software packages are available from their respective distribution channels.

- WGCNA, as presented in [56] and used in this study, is available in the Zenodo repository.
- wTO and CoDiNA packages are available on CRAN (https://cran.r-project.org/web/packages/wTO/index.html,
- https://cran.r-project.org/web/packages/CoDiNA/index.html) and the codes are available on GitHub (https://github.com/cran/wTO, https://github.com/cran/CoDiNA).
- PathVisio is available on https://pathvisio.org/downloads and Cytoscape is available on https://cytoscape.org/. CyTargetLinker is a Cytoscape App that can be installed and used through Cytoscape user interface. PathVisio/Cytoscape/CyTargetLinker workflow description is available in the Zenodo repository.
- EnrichNet is available on https://enrichnet.org/. EnrichNet code is available in the Zenodo repository to run locally with custom datasets.
- pathfindR is available on CRAN (https://cran.r-project.org/web/packages/pathfindR/index.html) and the code is available on GitHub (https://github.com/egeulgen/pathfindR).
- MOGAMUN is available on Bioconductor (https://bioconductor.org/packages/release/bioc/html/MOGAMUN.html) and the code is available on GitHub (https://github.com/elvanov/MOGAMUN).
- orsum is available on https://anaconda.org/bioconda/orsum and the codes are available on https://github.com/ozanozisik/orsum.

## Competing interests

The authors declare that they have no competing interests.

## Funding

This study has received funding from the European Union’s Horizon 2020 research and innovation programme under the EJP RD COFUND-EJP N° 825575. OO received funding from the Excellence Initiative of Aix-Marseille University - A*Midex, a French “Investissements d’Avenir programme”, and from the « Priority Research Programme on Rare Diseases » of the French Investments for the Future Programme. NSK received funding from the Scientific and Technical Research Council of Türkiye (TUBITAK) and from the PRIMUS Research Programme from Charles University. TA-D, MT, NQ-R, AJ, FE, EM received funding from European Union’s Horizon 2020 research and innovation programme under the EJP RD COFUND-EJP N° 825575.

## Authors’ contributions

OO: Data Curation, Investigation, Writing – Original Draft Preparation, Writing – Review & Editing; NSK: Investigation, Writing – Original Draft Preparation; TA-D: Investigation, Writing – Original Draft Preparation; MT: Investigation; NQ-R:Writing – Original Draft Preparation; AJ:Writing – Original Draft Preparation; OUS: Funding Acquisition, Supervision; MR: Funding Acquisition, Supervision; CTE: Funding Acquisition, Supervision; AB: Funding Acquisition, Supervision, Writing – Original Draft Preparation; FE: Conceptualization, Writing – Original Draft Preparation; EM: Conceptualization, Investigation, Project Administration, Writing – Original Draft Preparation, Writing – Review & Editing. All authors read and approved the final manuscript.

## Acknowledgements

We thank the members of the European Joint Programme on Rare Diseases.

## Ethics approval and consent to participate

Not applicable.

## Consent for publication

Not applicable.

## Notes

https://doi.org/10.5281/zenodo.8174258

### Competing Interest Statement

The authors have declared no competing interest.

